# KDM5C is a sex-biased brake against germline gene expression programs in somatic lineages

**DOI:** 10.1101/2024.11.08.622665

**Authors:** Katherine M. Bonefas, Ilakkiya Venkatachalam, Shigeki Iwase

**Author notes:** Correspondence should be addressed to K. Bonefas and S. Iwase.

## Abstract

The division of labor among cellular lineages is a pivotal step in the evolution of multicellularity. In mammals, the soma-germline boundary is formed during early embryogenesis, when genes that drive germline identity are repressed in somatic lineages through DNA and histone modifications at promoter CpG islands (CGIs). Somatic misexpression of germline genes is a signature of cancer and observed in select neurodevelopmental disorders. However, it is currently unclear if all germline genes use the same repressive mechanisms and if factors like development and sex influence their dysregulation. Here, we examine how cellular context influences the formation of somatic tissue identity in mice lacking lysine demethylase 5c (KDM5C), an X chromosome eraser of histone 3 lysine 4 di and tri-methylation (H3K4me2/3). We found male *Kdm5c* knockout (-KO) mice aberrantly express many tissue-specific genes within the brain, the majority of which are unique to the germline. By developing a comprehensive list of mouse germline-enriched genes, we observed *Kdm5c*-KO cells aberrantly express key drivers of germline fate during early embryogenesis but late-stage spermatogenesis genes within the mature brain. KDM5C binds CGIs within germline gene promoters to facilitate DNA CpG methylation as embryonic stem cells differentiate into epiblast-like cells (EpiLCs). However, the majority of late-stage spermatogenesis genes expressed within the *Kdm5c*-KO brain did not harbor promoter CGIs. These CGI-free germline genes were not bound by KDM5C and instead expressed through ectopic activation by RFX transcription factors. Furthermore, germline gene repression is sexually dimorphic, as female EpiLCs require a higher dose of KDM5C to maintain germline silencing. Altogether, these data revealed distinct regulatory classes of germline genes and sex-biased silencing mechanisms in somatic cells.

## Introduction

The separation of germline and somatic cellular identity is a pivotal step in the evolution of multicellularity and sexual reproduction^1–4^. In mammals, chromatin regulators decommission germline genes in somatic lineages when the early embryo transitions from naïve to primed pluripotency. Germline gene promoters initially gain repressive histone H2A lysine 119 monoubiquitination (H2AK119ub1)^5^ and histone H3 lysine 9 trimethylation (H3K9me3)^5,6^ in embryonic stem cells (ESCs) and are then decorated with DNA CpG methylation (CpGme) at their CpG islands (CGIs) in post-implantation epiblast cells^6–9^. While the silencing mechanisms for genes that establish germline identity are well characterized, it is unclear if other types of germline genes employ the same silencing mechanisms, such as those involved in the later stages of oogenesis and spermatogenesis. Furthermore, because many studies have focused on the silencing of key marker genes during early male embryonic development, much is unknown about how cellular context (i.e. sex and tissue environment) influences the manifestation of germline gene misexpression. Intriguingly, impaired soma-germline demarcation is a signature of aggressive cancers and observed in select neurodevelopmental disorders (NDDs)^10–13^. Thus, elucidating how cell context contributes to germline gene dysregulation will reveal novel mechanisms governing these pathologies.

Here, we employed genome-wide analyses to explore the loss of tissue identity in mice lacking the chromatin regulator lysine demethylase 5C (KDM5C, also known as SMCX or JARID1C). KDM5C lies on the X chromosome and erases histone 3 lysine 4 di-and trimethylation (H3K4me2/3), a permissive chromatin modification enriched at gene promoters^14^. Somatic loss of KDM5C promotes tumorigenicity in a variety of cancer types^15–17^, while pathogenic germline mutations cause the NDD Intellectual Developmental Disorder, X-linked, Syndromic, Claes-Jensen Type (MRXSCJ, OMIM: 300534). MRXSCJ is more common and severe in males and its neurological phenotypes include intellectual disability, seizures, aberrant aggression, and autistic behaviors^18–20^. Male *Kdm5c* knockout (-KO) mice recapitulate key MRXSCJ phenotypes, including hyperaggression, increased seizure propensity, social deficits, and learning impairments^21–23^. RNA sequencing (RNA-seq) of the *Kdm5c*-KO hippocampus revealed ectopic expression of some testis germline genes within the brain^22^. However, it is unclear if other tissue-specific genes are aberrantly transcribed with KDM5C loss, at what point in development germline gene misexpression begins, what mechanisms underlie their dysregulation, and how KDM5C interacts with other known germline silencing mechanisms.

To illuminate KDM5C’s role in tissue identity, we characterized the aberrant expression of tissue-enriched genes within the *Kdm5c*-KO brain and epiblast-like stem cells (EpiLCs), an *in vitro* model of the post-implantation embryo. We curated a list of mouse germline-enriched genes, enabling genome-wide analysis of germline gene silencing mechanisms for the first time. We identified two classes of germline genes based on their promoter CpG island content, which are dysregulated with KDM5C loss by distinct mechanisms and in a sex-biased manner.

## Results

### Tissue-enriched genes are aberrantly expressed in the *Kdm5c*-KO brain

Previous RNA sequencing (RNA-seq) of the adult male *Kdm5c*-KO hippocampus revealed ectopic expression of some germline genes unique to the testis^22^. It is currently unknown if the testis is the only tissue type misexpressed in the *Kdm5c*-KO brain. We first systematically tested whether other tissue-specific genes are misexpressed in the male brain with constitutive knockout of *Kdm5c* (*Kdm5c^-/y^*, 5CKO in figures)^24^ by using a published list of mouse tissue-enriched genes^25^.

We found a large proportion of significantly upregulated genes (DESeq2^26^, log2 fold change > 0.5, q < 0.1) within the male *Kdm5c*-KO amygdala and hippocampus are non-brain, tissue-specific genes (Amygdala: 0/0 up DEGs, NaN%; Hippocampus: 0/0 up DEGs, NaN%) (Figure 1A-B, Supplementary Table 1). For both the amygdala and hippocampus, the majority of tissue-enriched differentially expressed genes (DEGs) were testis genes (Figure 1A-B). Even though the testis has the largest total number of tissue-enriched genes (2,496 genes) compared to any other tissue, testis-enriched DEGs were significantly enriched in both brain regions (Amygdala p = 1.83e-05, Odds Ratio = 5.13; Hippocampus p = 4.26e-11, Odds Ratio = 4.45, Fisher’s Exact Test). An example of a testis-enriched gene misexpressed in the *Kdm5c*-KO brain is *FK506 binding protein 6 (Fkbp6)*, a known regulator of PIWI-interacting RNAs (piRNAs) and meiosis^27,28^ (Figure 1C). Interestingly, we also observed significant enrichment of ovary-enriched genes in both the amygdala and hippocampus (Amygdala p = 0.00574, Odds Ratio = 18.7; Hippocampus p = 0.048, Odds Ratio = 5.88, Fisher’s Exact Test) (Figure 1A-B). Ovary-enriched DEGs included *Zygotic arrest 1 (Zar1)*, which sequesters mRNAs in oocytes for meiotic maturation^29^ (Figure 1D). Given that the *Kdm5c*-KO mice we analyzed are male, these data demonstrate that the ectopic expression of gonad-enriched genes is independent of organismal sex.

**Figure 1:**
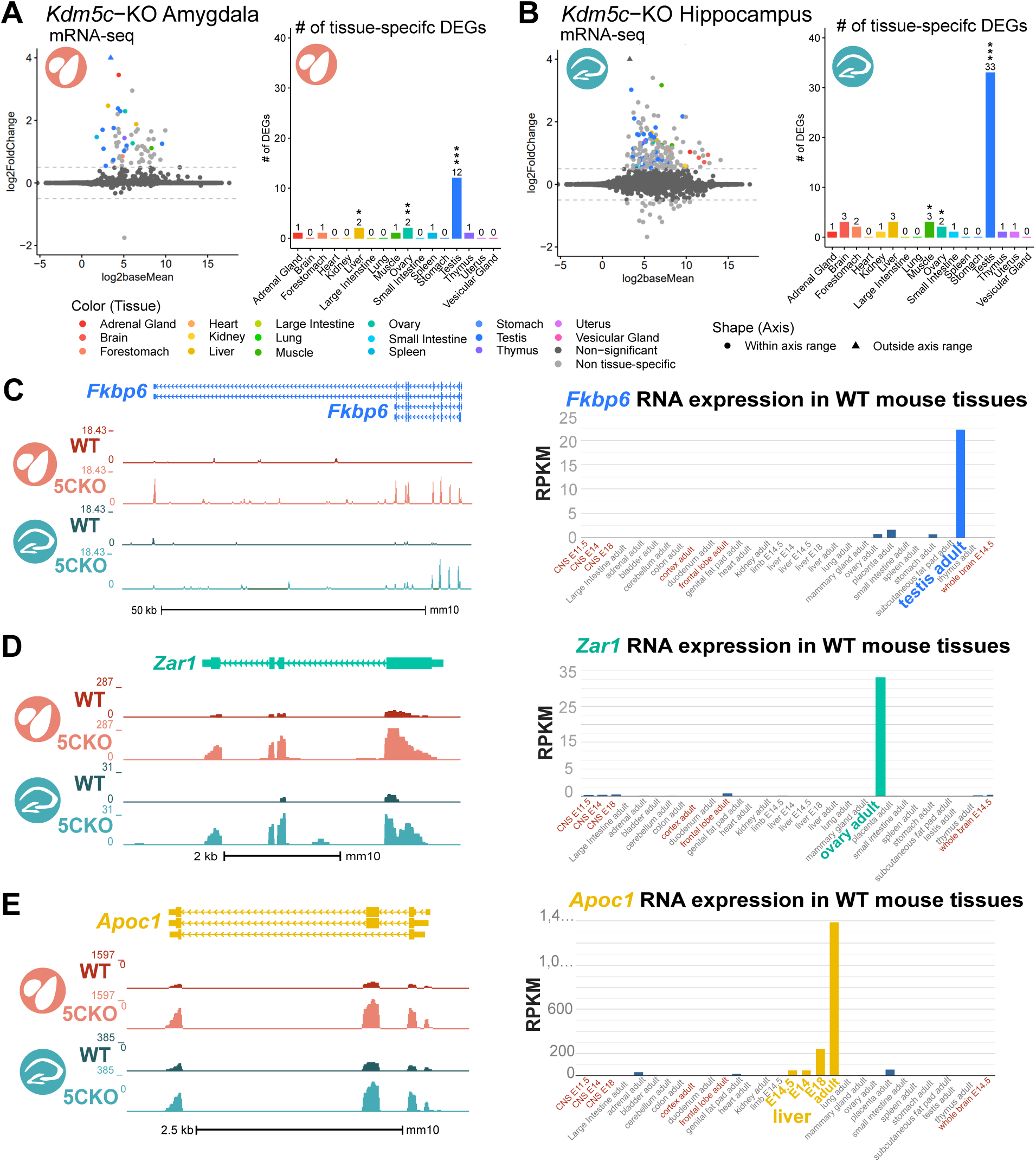
Tissue-enriched genes are misexpressed in the *Kdm5c*-KO brain. **A-B.** Expression of tissue-enriched genes (Li et al 2017) in the male *Kdm5c*-KO amygdala (A) and hippocampus (B). Left -MA plot of mRNA-sequencing. Right - Number of tissue-enriched differentially expressed genes (DEGs). * p<0.05, ** p<0.01, *** p<0.001, Fisher’s Exact Test **C.** Left - UCSC browser view of an example aberrantly expressed testis-enriched DEG, *FK506 binding protein 6 (Fkbp6)* in the wild-type (WT) and *Kdm5c*-KO (5CKO) amygdala (red) and hippocampus (teal) (Average, n = 4). Right - Expression of *Cyct* in wild-type tissues from NCBI Gene, with testis highlighted in blue and brain tissues highlighted in red. **D.** Left - UCSC browser view of an example ovary-enriched DEG, *Zygotic arrest 1 (Zar1)*. Right - Expression of *Zar1* in wild-type tissues from NCBI Gene, with ovary highlighted in teal and brain tissues highlighted in red. **E.** Left - UCSC browser view of an example liver-enriched DEG, *Apolipoprotein C-I (Apoc1)*. Right - Expression of *Apoc1* in wild-type tissues from NCBI Gene, with liver highlighted in orange and brain tissues highlighted in red.

Although not consistent across brain regions, we also found significant enrichment of genes biased towards two non-gonadal tissues - the liver (Amygdala p = 0.04, Odds Ratio = 6.58, Fisher’s Exact Test) and muscle (Hippocampus p = 0.01, Odds Ratio = 6.95, Fisher’s Exact Test) (Figure 1A-B). These include *Apolipoprotein C-I (Apoc1)*, a lipoprotein metabolism and transport gene^30^ (Figure 1E,see Discussion).

Our analysis of oligo(dT)-primed libraries^24^ indicates aberrantly expressed mRNAs are polyadenylated and spliced into mature transcripts in the *Kdm5c*-KO brain (Figure 1C-E). Of note, we observed little to no dysregulation of brain-enriched genes (Amygdala p = 1, Odds Ratio = 1.22; Hippocampus p = 0.74, Odds Ratio = 1.22, Fisher’s Exact Test), despite the fact these are brain samples and the brain has the second highest total number of tissue-enriched genes (708 genes). Altogether, these results suggest the aberrant expression of tissue-enriched genes within the brain is a major effect of KDM5C loss.

### Germline genes are misexpressed in the *Kdm5c*-KO brain

*Kdm5c*-KO brain expresses testicular germline genes^22^ (Figure 1), however the testis also contains somatic cells that support hormone production and germline functions. To determine if *Kdm5c*-KO results in ectopic expression of testicular somatic genes, we first evaluated the known functions of testicular DEGs through gene onotology. We found *Kdm5c*-KO testis-enriched DEGs had high enrichment of germline- relevant ontologies, including spermatid development (GO: 0007286, p.adjust = 6.2e-12) and sperm axoneme assembly (GO: 0007288, p.adjust = 2.45e-14) (Figure 2A, Supplementary Table 1).

**Figure 2:**
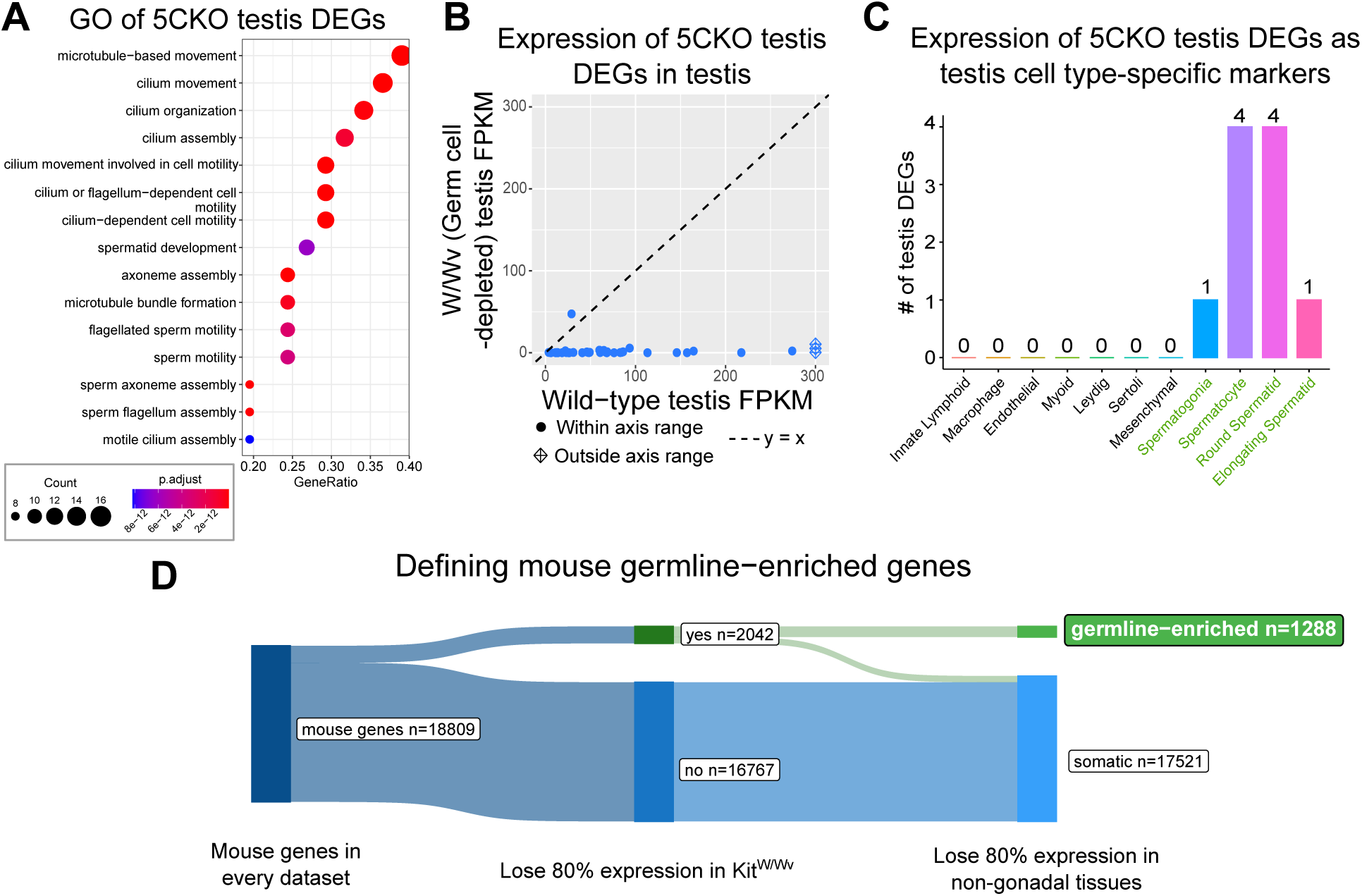
Aberrant transcription of germline genes in the *Kdm5c*-KO in the brain. **A.** enrichPlot gene ontology (GO) of *Kdm5c*-KO amygdala and hippocampus testis-enriched DEGs **B.** Expression of testis DEGs in wild-type (WT) testis versus germ cell-depleted (W/Wv) testis (Mueller et al 2013). Expression is in Fragments Per Kilobase of transcript per Million mapped reads (FPKM). **C.** Number of testis DEGs that were classified as cell-type specific markers in a single cell RNA-seq dataset of the testis (Green et al 2018). Germline cell types are highlighted in green, somatic cell types in black. **D.** Sankey diagram of mouse genes filtered for germline enrichment based on their expression in wild-type and W/Wv mice and in adult mouse non-gonadal tissues (Li et al 2017).

We then evaluated *Kdm5C*-KO testicular DEG expression in wild-type testes versus testes with germ cell depletion^31^, which was accomplished by heterozygous *W* and *Wv* mutations in the enzymatic domain of *c-Kit* (Kit^W/Wv^)^32^. Almost all *Kdm5c*-KO testis-enriched DEGs lost expression with germ cell depletion (Figure 2B). We then assessed testis-enriched DEG expression in a published single cell RNA-seq dataset that identified cell type-specific markers within the testis^33^. Some *Kdm5c*-KO testis-enriched DEGs were classified as specific markers for different germ cell developmental stages (e.g. spermatogonia, spermatocytes, round spermatids, and elongating spermatids), yet none marked somatic cells (Figure 2C). Together, these data demonstrate that the *Kdm5c*-KO brain aberrantly expresses germline genes but not somatic testicular genes, reflecting an erosion of the soma-germline boundary.

As of yet, research on germline gene silencing mechanisms has focused on a handful of key genes rather than assessing germline gene suppression genome-wide, due to the lack of a comprehensive gene list. We therefore generated a list of mouse germline-enriched genes using RNA-seq datasets of Kit^W/Wv^ mice that included males and females at embryonic day 12, 14, and 16^34^ and adult male testes^31^. We defined genes as germline-enriched if their expression met the following criteria: 1) their expression is greater than 1 FPKM in wild-type gonads 2) their expression in any non-gonadal tissue of adult wild type mice^25^ does not exceed 20% of their maximum expression in the wild-type germline, and 3) their expression in the germ cell-depleted gonads, for any sex or time point, does not exceed 20% of their maximum expression in the wild-type germline. These criteria yielded 1,288 germline-enriched genes (Figure 2D), which was hereafter used as a resource to globally characterize germline gene misexpression with *Kdm5c* loss (Supplementary Table 2).

### *Kdm5c*-KO epiblast-like cells aberrantly express key regulators of germline identity

Germ cells are typically distinguished from somatic cells soon after the embryo implants into the uterine wall^35,36^, when germline genes are silenced in epiblast stem cells that will form the somatic tissues^37^. This developmental time point can be modeled *in vitro* through differentiation of naïve embryonic stem cells (nESCs) into epiblast-like stem cells (EpiLCs) (Figure 3A)^38,39^. While some germline-enriched genes are also expressed in nESCs and in the 2-cell stage^40–42^, they are silenced as they differentiate into EpiLCs^6,7^. Therefore, we tested if KDM5C was necessary for the initial silencing of germline genes in somatic lineages by evaluating the impact of *Kdm5c* loss in male EpiLCs.

**Figure 3:**
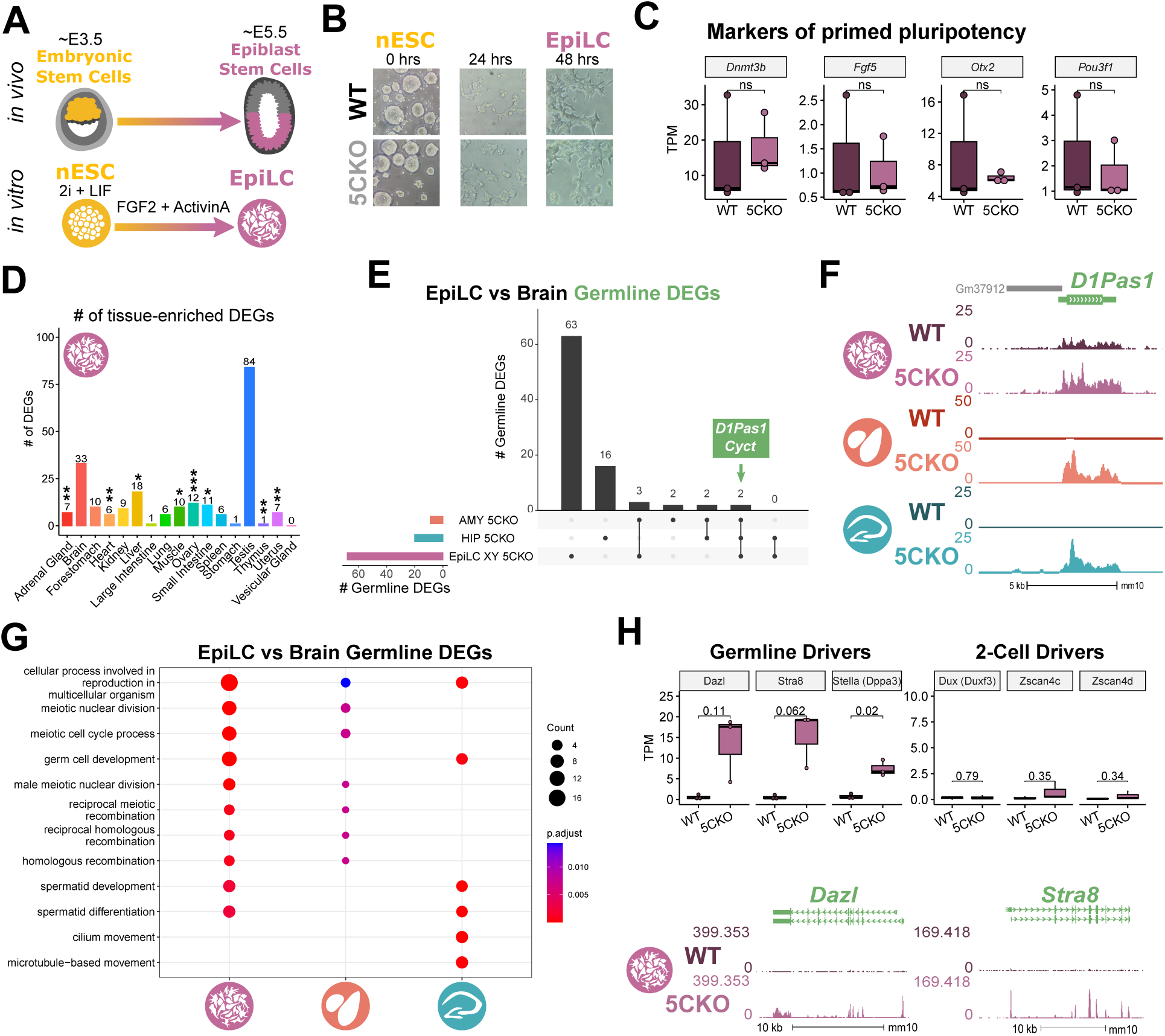
*Kdm5c*-KO epiblast-like cells express key drivers of germline identity. **A.** Top - Diagram of *in vivo* differentiation of embryonic stem cells (ESCs) of the inner cell mass into epiblast stem cells. Bottom - *in vitro* differentiation of ESCs into epiblast-like cells (EpiLCs). **B.** Representative images of male wild-type (WT) and *Kdm5c*-KO (5CKO) cells during ESC to EpiLC differentiation. Brightfield images taken at 20X. **C.** No significant difference in primed pluripotency maker expression in wild-type versus *Kdm5c*-KO EpiLCs. Welch’s t-test, expression in transcripts per million (TPM). **D.** Number of tissue-enriched differentially expressed genes (DEGs) in *Kdm5c*-KO EpiLCs. * p<0.05, ** p<0.01, *** p<0.001, Fisher’s Exact Test. **E.** Upset plot displaying the overlap of germline DEGs expressed in *Kdm5c*-KO EpiLCs, amygdala (AMY), and hippocampus (HIP) RNA-seq datasets. **F.** UCSC browser view of an example germline gene, *D1Pas1*, that is dysregulated *Kdm5c*-KO EpiLCs (top, purple. Average, n = 3), amygdala (middle, red. Average, n = 4), and hippocampus (bottom, blue. Average, n = 4). **G.** enrichPlot gene ontology analysis comparing enriched biological processes for *Kdm5c*-KO EpiLC, amygdala, and hippocampus germline DEGs. **H.** Top left - Example germline identity DEGs unique to EpiLCs. Top right - Example 2-cell genes that are not dysregulated in *Kdm5c*-KO EpiLCs. p-values for Welch’s t-test. Bottom - UCSC browser view of *Dazl* and *Stra8* expression in wild-type and *Kdm5c*-KO EpiLCs (Average, n = 3).

*Kdm5c*-KO cell morphology during ESC to EpiLC differentiation appeared normal (Figure 3B) and EpiLCs properly expressed markers of primed pluripotency, such as *Dnmt3b*, *Fgf5*, *Pou3f1*, and *Otx2* (Figure 3C). We then identified tissue-enriched DEGs in a RNA-seq dataset of wild-type and *Kdm5c*-KO EpiLCs^43^ (DESeq2, log2 fold change > 0.5, q < 0.1, Supplementary Table 3). Similar to the *Kdm5c*-KO brain, we observed general dysregulation of tissue-enriched genes, with the largest number of genes belonging to the brain and testis, although they were not significantly enriched (Figure 3D). Using our list of mouse germline-enriched genes assembled above, we identified 68 germline genes misexpressed in male *Kdm5c*-KO EpiLCs.

We then compared EpiLC germline DEGs to those expressed in the *Kdm5c*-KO brain to determine if germline genes are constitutively dysregulated or change over the course of development. The majority of germline DEGs were unique to either EpiLCs or the brain, with only *D1Pas1* and *Cyct* shared across all tissue/cell types (Figure 3E-F). EpiLC germline DEGs had particularly high enrichment of meiosis-related gene ontologies when compared to the brain (Figure 3G, Supplementary Table 3), such as meiotic cell cycle process (GO:1903046, p.adjust = 2.2e-07) and meiotic nuclear division (GO:0140013, p.adjust = 1.37e-07). While there was modest enrichment of meiotic gene ontologies in both brain regions, the *Kdm5c*-KO hippocampus primarily expressed late-stage spermatogenesis genes involved in sperm axoneme assembly (GO:0007288, p.adjust = 0.00621) and sperm motility (GO:0097722, p.adjust = 0.00612).

Notably, DEGs unique to *Kdm5c*-KO EpiLCs included key drivers of germline identity, such as *Stimulated by retinoic acid 8* (*Stra8*: log2 fold change = 3.73, q = 2.17e-39) and *Deleted in azoospermia like* (*Dazl*: log2 fold change = 3.36, q = 3.19e-12) (Figure 3H). These genes are typically expressed when a subset of epiblast stem cells become primordial germ cells (PGCs) and then again in mature germ cells to trigger meiotic gene expression programs^44–46^. Of note, some germline genes, including *Dazl*, are also expressed in the two-cell embryo^41,47^. However, we did not see derepression of two-cell stage-specific genes, like *Duxf3 (Dux)* (log2 fold change = -0.282, q = 0.337) and *Zscan4d* (log2 fold change = 0.25, q = 0.381) (Figure 3H, Supplementary Table 3), indicating *Kdm5c*-KO EpiLCs do not revert back to a 2-cell state. Altogether, *Kdm5c*-KO EpiLCs express key drivers of germline identity and meiosis while the brain primarily expresses spermiogenesis genes, indicating germline gene misexpression mirrors germline development during the progression of somatic development.

### Female epiblast-like cells have heightened germline gene misexpression with *Kdm5c* loss

It is currently unknown if the misexpression of germline genes is influenced by sex, as previous studies on germline gene repressors have focused on male cells^5,6,8,48,49^. Sex is particularly pertinent in the case of KDM5C because it partially escapes X chromsome inactivation (XCI), resulting in a higher dosage in females^50–53^. We therefore explored the impact of chromosomal sex upon germline gene suppression by comparing their dysregulation in male *Kdm5c* hemizygous knockout (*Kdm5c^-/y^*, XY *Kdm5c*-KO, XY 5CKO), female homozygous knockout (*Kdm5c^-/-^*, XX *Kdm5c*-KO, XX 5CKO), and female heterozygous knockout (*Kdm5c^-/+^*, XX *Kdm5c*-HET, XX 5CHET) EpiLCs^43^.

In EpiLCs, homozygous and heterozygous *Kdm5c* knockout females expressed over double the number of germline-enriched genes than hemizygous males (Figure 4A, Supplementary Table 3). While the majority of germline DEGs in *Kdm5c*-KO males were also dysregulated in females (74%), many were sex-specific, such as *Tktl2* and *Esx1* (Figure 4B). We then compared the known functions of germline genes dysregulated uniquely in males and females or misexpressed in all samples (Figure 4C, Supplementary Table 3). Female-specific germline DEGs were enriched for meiotic (GO:0051321 - meiotic cell cycle, p.adjust = 7.81E-14) and flagellar (GO:0003341 - cilium movement, p.adjust = 4.87E-06) functions, while male-specifc DEGs had roles in mitochondrial and cell signaling (GO:0070585 - protein localization to mitochondrion, p.adjust = 0.025).

**Figure 4:**
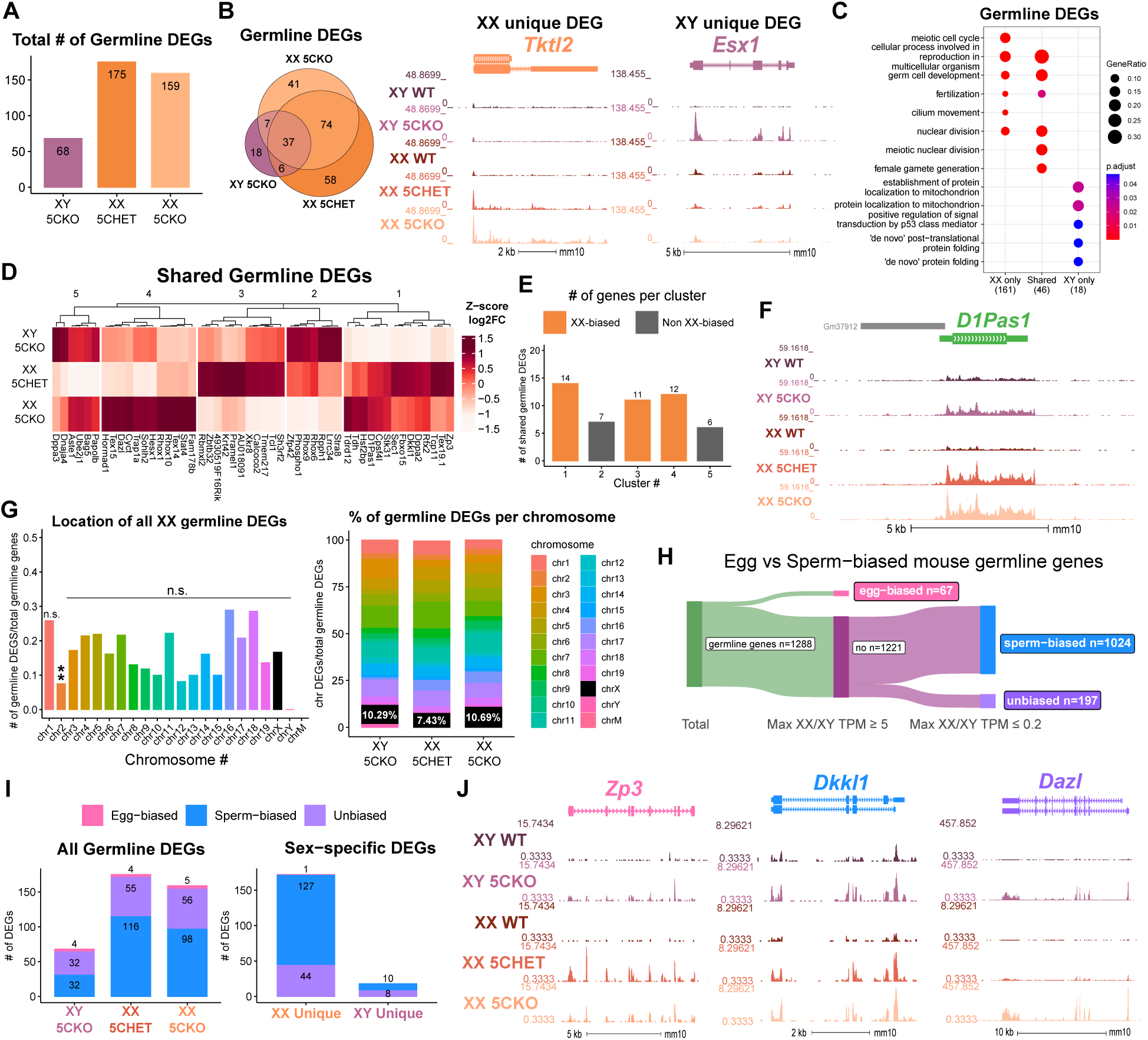
Chromosomal sex influences *Kdm5c*-KO germline gene misexpression. **A.** Total number of germline-enriched RNA-seq DEGs for male hemizygous *Kdm5c* knockout EpiLCs (XY 5CKO, purple), female heterozygous *Kdm5c* knockout (XX 5CHET, orange), female homozygous *Kdm5c* knockout (XX 5CKO, light orange) EpiLCs. **B.** Left - Eulerr overlap of *Kdm5c* mutant male and female EpiLC germline DEGs. Right - Example of germline DEGs unique to females or males, *Tktl2* and *Esx1*.**C.** enrichPlot gene ontology analysis comparing enriched biological processes for germline DEGs shared between *Kdm5c* mutant males and females (Shared), or unique to one sex (XX only or XY only). **D.** Heatmap of germline DEGs shared between male and female mutants. Color is the average log 2 fold change from sex-matched wild-type, z-scored across rows. **E.** Number of genes within each cluster from D. Clusters with higher expression in females compared to males (XX-biased) highlighted in orange. **F.** UCSC browser view of a male and female shared germline DEG *D1Pas1* that is more highly expressed in female mutants (Average, n = 3). **G.** Left - Number of all female germline DEGs located on each chromosome over the total number of germline-enriched genes on that chromsome. P-values for Fisher Exact Test, ** p < 0.01, n.s. non-significant. Germline DEGs were only significant for chromosome 2, in which they were significantly depleted. Right - Percentage of germline DEGs that lie on each chromosome for each *Kdm5c* mutant. X chromosome highlighted in black. **H.** Sankey diagram classifying egg-biased (pink) and sperm-biased (blue) and unbiased (purple) mouse germline-enriched genes. **I.** Number of egg, sperm, or unbiased germline DEGs for male and female *Kdm5c* mutants. **J.** UCSC browser view of egg-biased (*Zp3*), sperm-biased (*Dkkl1*), and unbiased (*Dazl*) germline genes dysregulated in both male and female *Kdm5c* mutants (Average of n = 3).

The majority of germline genes expressed in both sexes were more highly dysregulated in females compared to males (Figure 4D-F). This increased degree of dysregulation in females, along with the increased total number of germline genes, indicates females are more sensitive to losing KDM5C-mediated germline gene suppression. Heightened germline gene dysregulation in females could be due to impaired XCI in *Kdm5c* mutants^43^, as many spermatogenesis genes lie on the X chromosome^54,55^. However, female germline DEGs were not biased towards the X chromosome (p = 1, Odds Ratio = 0.96, Fisher’s Exact Test) and females had a a similar overall proportion of germline DEGs belonging to the X chromosome as males (XY *Kdm5c*-KO - 10.29%, XX *Kdm5c*-HET - 7.43%, XX *Kdm5c*-KO - 10.59%) (Figure 4G). The majority of germline DEGs instead lie on autosomes for both male and female *Kdm5c* mutants (Figure 4G). Thus, while female EpiLCs are more prone to germline gene misexpression with KDM5C loss, it is likely independent of XCI defects.

### Germline gene misexpression in *Kdm5c* mutants is independent of germ cell sex

Although many germline genes have shared functions in the male and female germline, e.g. PGC formation, meiosis, and genome defense, some have unique or sex-biased expression. Therefore, we wondered if *Kdm5c* mutant males would primarily express sperm genes while mutant females would primarily express egg genes. To comprehensively assess whether germline gene sex corresponds with *Kdm5c* mutant sex, we first filtered our list of germline-enriched genes for egg and sperm-biased genes (Figure 4, Supplementary Table 2). We defined germ cell sex-biased genes as those whose expression in the opposite sex, at any time point, is no greater than 20% of the gene’s maximum expression in a given sex. This criteria yielded 67 egg-biased, 1,024 sperm-biased, and 197 unbiased germline-enriched genes. We found regardless of sex, egg, sperm, and unbiased germline genes were dyregulated in all *Kdm5c* mutants at similar proportions (Figure 4I-J). Furthermore, germline genes dysregulated exclusively in either male or female mutants were also not biased towards their corresponding germ cell sex (Figure 4I). Altogether, these results demonstrate sex differences in germline gene dysregulation is not due to sex-specific activation of sperm or egg transcriptional programs.

### KDM5C binds to a subset of germline gene promoters during early embryogenesis

KDM5C binds to the promoters of several germline genes in embryonic stem cells (ESCs) but not in neurons^22,56^. However, due to the lack of a comprehensive list of germline-enriched genes, it is unclear if KDM5C is enriched at germline gene promoters, what types of germline genes KDM5C regulates, and if its binding is maintained at any germline genes in neurons.

To address these questions, we analyzed KDM5C chromatin immunoprecipitation followed by DNA sequencing (ChIP-seq) datasets in EpiLCs^43^ and primary forebrain neuron cultures (PNCs)^21^ (MACS2 q < 0.1, fold enrichment > 1, and removal of false-positive *Kdm5c*-KO peaks). EpiLCs had a higher total number of high-confidence KDM5C peaks than PNCs (EpiLCs: 5,808, PNCs: 1,276). KDM5C was primarily localized to gene promoters in both cell types (promoters = transcription start site (TSS) ± 500 bp, EpiLCs: 4,190, PNCs: 745), although PNCs showed increased localization to non-promoter regions (Figure 5A).

**Figure 5:**
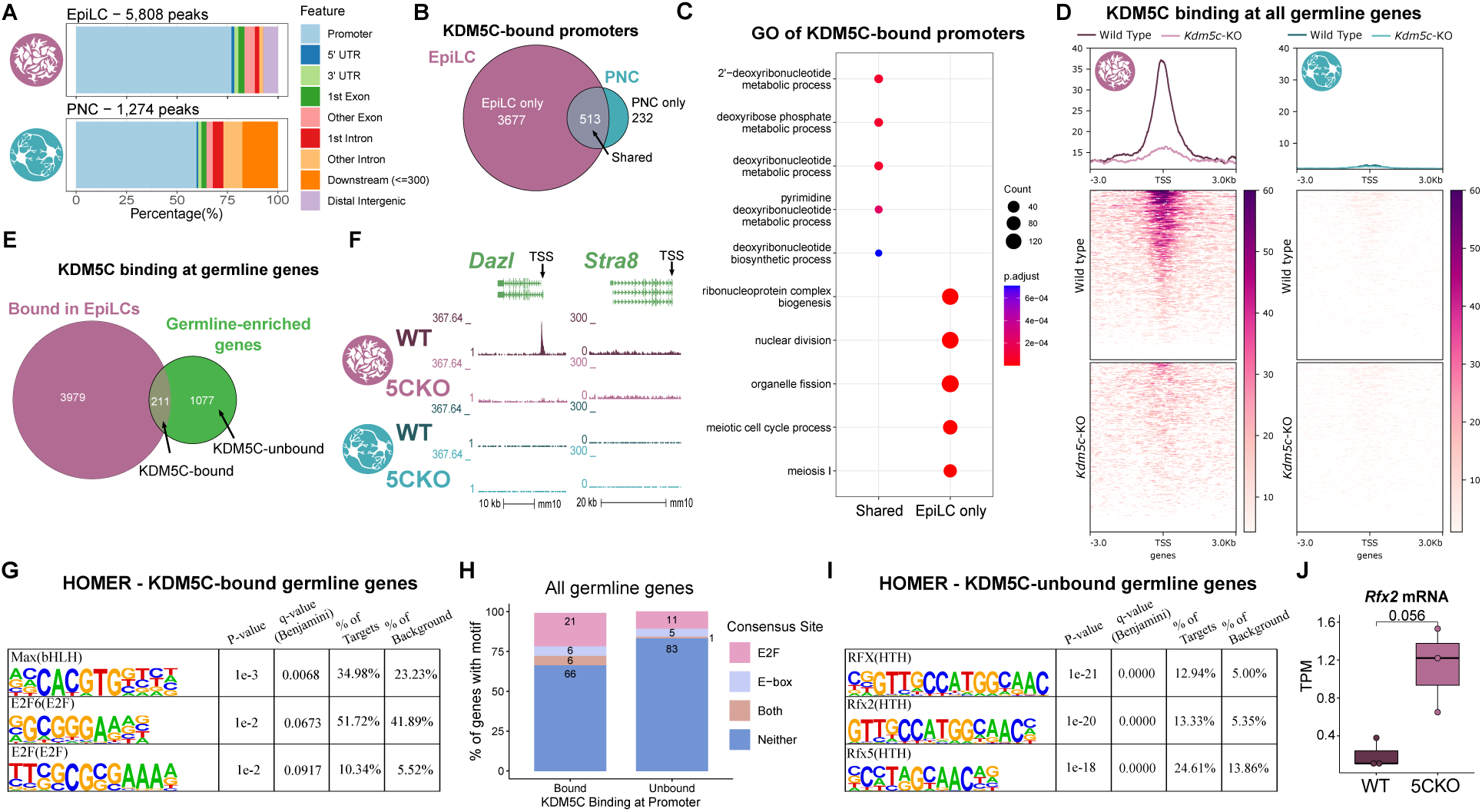
KDM5C binds to a subset of germline gene promoters during early embryogenesis. **A.** ChIPseeker localization of KDM5C peaks at different genomic regions in EpiLCs (top) and hippocampal and cortex primary neuron cultures (PNCs, bottom). **B.** Overlap of genes with KDM5C bound to their promoters (TSS *±* 500) in EpiLCs (purple) and PNCs (blue). **C.** Gene ontology (GO) comparison of genes with KDM5C bound to their promoter in EpiLCs and PNCs. Genes were classified as either bound in EpiLCs only (EpiLC only), unique to PNCs (PNC only, no significant ontologies) or bound in both PNCs and EpiLCs (shared). **D.** Average KDM5C binding around the transcription start site (TSS) of all germline-enriched genes in EpiLCs (left) and PNCs (right). **E.** Eulerr overlap of germline-enriched genes (green) with significant KDM5C binding at their promoter in EpiLCs (purple). **F.** Example KDM5C ChIP-seq signal around the *Dazl* TSS but not *Stra8* in EpiLCs. **G.** HOMER motif analysis of all KDM5C-bound germline gene promoters, highlighting significant enrichment of MAX, E2F6, and E2F motifs. **H.** Number of all gene promoters bound or unbound by KDM5C with instances of the E2F or E-box consensus sequence. **I.** HOMER motif analysis of all KDM5C-unbound germline gene promoters, highlighting significant enrichment of RFX family transcription factor motifs. **J.** Expression of RNA-seq DEG *Rfx2* in wild-type and *Kdm5c*-KO EpiLCs. P-value of Welch’s t-test, expression in transcripts per million (TPM).

**Figure 6:**
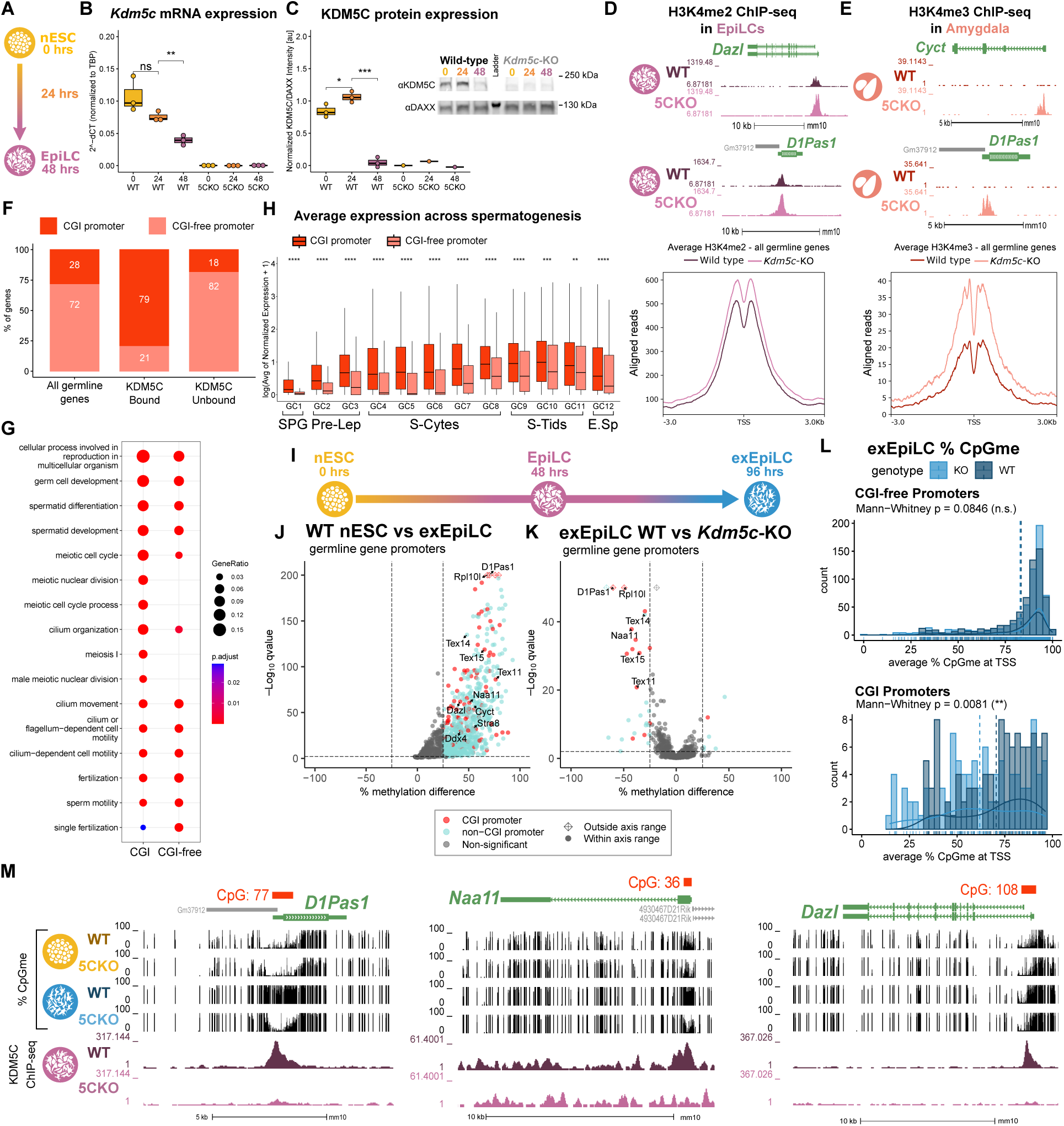
KDM5C promotes long-term silencing of germline genes via DNA methylation at CpG islands. **A.** Diagram of embryonic stem cell (ESC) to epiblast-like cell (EpiLC) differentiation and collection time points. **B.** Real time quantitative PCR (RT-qPCR) of *Kdm5c* mRNA expression in wild-type (WT) and *Kdm5c*-KO (5CKO) ESCs at 0, 24, and 48 hours of differentiation into EpiLCs. Expression calculated in comparison to TBP mRNA expression (2^-deltaCT^). * p < 0.05, ** p < 0.01, *** p < 0.001, Welch’s t-test. **C.** KDM5C protein expression normalized to DAXX. Quantified intensity using ImageJ (artificial units - au). Right - representative lanes of Western blot for KDM5C and DAXX. * p < 0.05, ** p < 0.01, *** p < 0.001, Welch’s t-test. **D.** Top - Representative UCSC browser view of histone 3 lysine 4 dimethylation (H3K4me2) ChIP-seq signal at two germline genes in wild-type and *Kdm5c*-KO EpiLCs. Bottom - Average H3K4me2 signal at the TSS of all germline-enriched genes in wild-type (dark purple) and *Kdm5c*-KO (light purple) EpiLCs. **E.** Top - Representative UCSC browser view of histone 3 lysine 4 trimethylation (H3K4me3) ChIP-seq signal at two germline genes in the wild-type (WT) and *Kdm5c*-KO (5CKO) adult amygdala. Bottom - Average H3K4me3 signal at the transcription start site (TSS) of all germline-enriched genes in wild-type (dark red) and *Kdm5c*-KO (light red) amygdala. **F.** Percentage of germline genes that harbor CpG islands (CGIs) in their promoters (CGI promoter, red), based on UCSC annotation. Left - percentage of all germline-enriched genes, middle - KDM5C-bound germline genes, and right - KDM5C-unbound germline genes. **G.** enrichPlot gene ontology analysis of germline genes with (CGI-promoter) or without (CGI-free) CGIs in their promoter. **H.** Expression of germline genes with (CGI-promoter, red) or without (CGI-free, salmon) CGIs in their promoter across stages of spermatogenesis from Green et al 2018. SPG - spermatogonia, Pre-Lep - preleptotene spermatocytes, S-Cytes - meiotic spermatocytes, S-Tids - post-meiotic haploid round spermatids, E.Sp - elongating spermatids. Wilcoxon test, * p < 0.05, ** p < 0.01, *** p < 0.001, **** p < 0.0001. **I.** Diagram of ESC to extended EpiLC (exEpiLC) differentiation. **J.** Volcano plot of whole genome bisulfite sequencing (WGBS) comparing CpG methylation (CpGme) at germline gene promoters (TSS *±* 500) in wild-type (WT) ESCs versus exEpiLCs. Significantly differentially methylated promoters (q < 0.01, |methylation difference| > 25%) with CGIs in red, CGI-free promoters in light blue **K.** Volcano plot of WGBS of wild-type (WT) versus *Kdm5c*-KO exEpiLCs for germline gene promoters. Promoters with CGIs in red, CGI-free promoters in light blue. **L.** Histogram of average percent CpGme at the promoter of germline genes with or without CGIs. Wild-type in navy and *Kdm5c*-KO (KO) in light blue. Dashed lines are average methylation for each genotype, p-values for Mann-Whitney U test. **M.** UCSC browser view of germline genes, showing UCSC annotated CGI with number of CpGs, representative CpGme in wild-type (WT) and *Kdm5c*-KO (5CKO) ESCs and exEpiLCs, and KDM5C ChIP-seq signal in EpiLCs.

The majority of promoters bound by KDM5C in PNCs were also bound in EpiLCs (513 shared promoters), however a large portion of gene promoters were bound by KDM5C only in EpiLCs (3,677 EpiLC only promoters) (Figure 5B). Genes bound by KDM5C in both PNCs and EpiLCs were enriched for functions involving nucleic acid turnover, such as deoxyribonucleotide metabolic process (GO:0009262, p.adjust = 8.28e-05) (Figure 5C, Supplementary Table 4). Germline ontologies were enriched only in EpiLC-specific, KDM5C-bound promoters, such as meiotic nuclear division (GO: 0007127 p.adjust = 6.77e-16) (Figure 5C). There were no significant ontologies for PNC-specific KDM5C target genes. Using our mouse germline gene list, we observed evident KDM5C signal around the TSS of many germline genes in EpiLCs, but not in PNCs (Figure 5D). Based on our ChIP-seq peak cut-off criteria, KDM5C was highly enriched at 211 germline gene promoters in EpiLCs (16.4% of all germline genes) (Figure 5E, Supplementary Table 2). Of note, KDM5C was only bound to about one third of RNA-seq DEG promoters unique to EpiLCs or the brain (EpiLC only DEGs: 34.9%, Brain only DEGs: 30%) (Supplementary Figure 1A-C). Representative examples of EpiLC DEGs bound and unbound by KDM5C in EpiLCs are *Dazl* and *Stra8*, repsectively (Figure 5F). However, the four of the five germline genes dysregulated in both EpiLCs and the brain were bound by KDM5C in EpiLCs (*D1Pas1*, *Hsf2bp*, *Cyct*, and *Stk31*) (Supplementary Figure 1A). Together, these results demonstrate KDM5C is recruited to a subset of germline genes in EpiLCs, including meiotic genes, but does not directly regulate germline genes in neurons. Furthermore, the majority of germline mRNAs expressed in *Kdm5c*-KO cells are dysregulated independent of direct KDM5C recruitment to their gene promoters, however genes dysregulated across *Kdm5c*-KO development are often direct KDM5C targets.

Many germline-specific genes are suppressed by the polycomb repressive complex 1.6 (PRC1.6), which contains the transcription factor heterodimers E2F6/DP1 and MGA/MAX that respectively bind E2F and E-box motifs within germline gene promoters^5,6,8,42,48,49,57–59^. PRC1.6 members may recruit KDM5C to germline gene promoters^22^, given their association with KDM5C in HeLa cells and ESCs^47,60^. We thus used HOMER^61^ to identify transcription factor motifs enriched at KDM5C-bound or unbound germline gene promoters (TSS ± 500 bp, q-value < 0.1, Supplementary Table 4). MAX and E2F6 binding sites were significantly enriched at germline genes bound by KDM5C in EpiLCs (MAX q-value: 0.0068, E2F6 q-value: 0.0673, E2F q-value: 0.0917), but not at germline genes unbound by KDM5C (Figure 5G). One third of KDM5C-bound promoters contained the consensus sequence for either E2F6 (E2F, 5’-TCCCGC-3’), MGA (E-box, 5’-CACGTG-3’), or both, but only 17% of KDM5C-unbound genes contained these motifs (Figure 5H). KDM5C-unbound germline genes were intstead enriched for multiple RFX transcription factor binding sites (RFX q-value < 0.0001, RFX2 q-value < 0.0001, RFX5 q-value < 0.0001) (Figure 5I, Supplementary figure 1D). RFX transcription factors bind X-box motifs^62^ to promote ciliogenesis^63,64^ and among them is RFX2, a central regulator of post-meiotic spermatogeneis^65,66^. Although *Rfx2* is also not a direct target of KDM5C (Supplementary Figure 1E), RFX2 mRNA is derepressed in *Kdm5c*-KO EpiLCs (Figure 5J). Thus, RFX2 is a candidate transcription factor for driving the ectopic expression of many KDM5C-unbound germline genes in *Kdm5c*-KO cells.

### KDM5C is recruited to CpG islands at germline promoters to facilitate *de novo* DNA methylation

Previous work found two germline gene promoters have a marked reduction in DNA CpG methylation (CpGme) in the adult *Kdm5c*-KO hippocampus^22^. Since histone H3K4me2/3 impede *de novo* CpGme^67,68^, KDM5C’s removal of H3K4me2/3 may be required to suppress germline genes. However, KDM5C’s catalytic activity was recently shown to be dispensible for suppressing *Dazl* in undifferentiated ESCs^47^. To reconcile these observations, we hypothesized KDM5C erases H3K4me2/3 to promote the initial placement of CpGme at germline gene promoters in EpiLCs.

To test this hypothesis, we first characterized KDM5C’s expression as naïve ESCs differentiate into EpiLCs (Figure 6A). While *Kdm5c* mRNA steadily decreased from 0 to 48 hours of diferentiation (Figure 6B), KDM5C protein initially increased from 0 to 24 hours and then decreased to near knockout levels by 48 hours (Figure 6C). We then characterized KDM5C’s substrates (H3K4me2/3) at germline gene promoters with *Kdm5c* loss using published ChIP-seq datasets^24,43^. *Kdm5c*-KO samples showed a marked increase in H3K4me2 in EpiLCs (Figure 6D) and H3K4me3 in the amygdala (Figure 6E) around the TSS of germline genes. Together, these data suggest KDM5C acts during the transition between ESCs and EpiLCs to remove H3K4me2/3 at germline gene promoters.

Germline genes accumulate CpG methylation (CpGme) at CpG islands (CGIs) during the transition from naïve to primed pluripotency^7,9,69^. We first examined how many of our germline-enriched genes had promoter CGIs (TSS ± 500 bp) using the UCSC genome browser^70^. Notably, out of 1,288 germline-enriched genes, only 356 (27.64%) had promoter CGIs (Figure 6F, Supplementary Table 2). CGI-containing germline genes had higher enrichment of meiotic gene ontologies compared to CGI-free genes, including meiotic nuclear division (GO:0140013, p.adjust = 2.17e-12) and meiosis I (GO:0007127, p.adjust = 3.91e-10) (Figure 6G, Supplementary Table 5). Germline genes with promoter CGIs were more highly expressed than CGI-free genes across spermatogenesis stages, with highest expression in meiotic spermatocytes (Figure 6H). Contrastingly, CGI-free genes only displayed substantial expression in post-meiotic round spermatids (Figure 6H). Although only a minor portion of germline gene promoters contained CGIs, CGIs strongly determined KDM5C’s recruitment to germline genes (p = 2.37e-67, Odds Ratio = 17.8, Fisher’s Exact Test), with 79.15% of KDM5C-bound germline gene promoters harboring CGIs (Figure 6F).

To assess how KDM5C loss impacts initial CpGme placement at germline gene promoters, we perfomed whole genome bisulfite sequencing (WGBS) in male wild-type and *Kdm5c*-KO ESCs and 96-hour extend EpiLCs (exEpiLCs), when germline genes reach peak methylation levels^6^ (Figure 6I). We first identified which germline gene promoters siginificantly gained CpGme in wild-type cells during nESC to exEpiLCs differentiation (methylKit^71^, q < 0.01, |methylation difference| > 25%, TSS ± 500 bp). In wild-type cells, the majority of germline genes gained substantial CpGme at their promoter during differentiation (60.08%), regardless if their promoter contained a CGI (Figure 6J, Supplementary Table 5).

We then identified promoters differentially methylated in wild-type versus *Kdm5c*-KO exEpiLCs (methylKit, q < 0.01, |methylation difference| > 25%, TSS ± 500 bp, Supplementary Table 5). Of the 48,882 promoters assessed, 274 promoters were significantly hypomethylated and 377 promoters were significantly hyper-methylated with KDM5C loss (Supplementary Figure 2A). Many promoters hyper-and hypomethylated in *Kdm5c*-KO exEpiLCs belonged to genes with unknown functions. However, 10.22% of hypomethy-lated promoters belonged to germline genes and germline-relevant ontologies like meiotic nuclear division (GO:0140013, p.adjust = 0.012) are significantly enriched (Supplementary Figure 2B, Supplementary Table 5). Approximately half of all germline gene promoters hypomethylated in *Kdm5c*-KO exEpiLCs are direct targets of KDM5C in EpiLCs (13 out of 28 hypomethylated promoters).

Promoters that showed the most robust loss of CpGme in *Kdm5c*-KO exEpiLCs (lowest q-values) harbored CGIs (Figure 6K). CGI promoters, but not CGI-free promoters, had a significant reduction in CpGme with KDM5C loss as a whole (Figure 6L) (Non-CGI promoters p = 0.0846, CGI promoters p = 0.0081, Mann-Whitney U test). Significantly hypomethylated promoters included germline genes consistently dysregulated across multiple *Kdm5c*-KO RNA-seq datasets^22^, such as *D1Pas1* (methlyation difference = -60.03%, q-value = 3.26e-153) and *Naa11* (methlyation difference = -42.45%, q-value = 1.44e-38) (Figure 6M). Unexpectedly, we observed only a modest reduction in CpGme at *Dazl*’s promoter (methlyation difference = -6.525%, q-value = 0.0159) (Figure 6N). Altogether, these results demonstrate KDM5C is recruited to germline gene CGIs in EpiLCs to promote CpGme at those promoters. Furthermore, our data suggest while KDM5C’s cataltyic activity is required for the repression of some germline genes, CpGme can be placed at others even with elevated H3K4me2/3 around the TSS.

## Discussion

In the above study, we demonstrate KDM5C’s pivotal role in the development of tissue identity. We first characterized tissue-enriched genes expressed within the mouse *Kdm5c*-KO brain and identified substantial derepression of testis, liver, muscle, and ovary-enriched genes. Testis genes significantly enriched within the *Kdm5c*-KO amygdala and hippocampus are specific to the germline and absent in somatic cells. *Kdm5c*-KO epiblast-like cells (EpiLCs) aberrantly express key drivers of germline identity and meiosis, including *Dazl* and *Stra8*, while the adult brain primarily expresses genes important for late spermatogenesis. We demonstrated that although sex did not influence whether sperm or egg-specific genes were misexpressed, female EpiLCs have heightened germline gene de-repression with KDM5C loss. Germline genes can become aberrantly expressed in *Kdm5c*-KO cells via indirect mechanisms, such as activation through ectopic RFX transcription factors. Finally, we found KDM5C is dynamically regulated during ESC to EpiLC differentiation to promote long-term germline gene silencing through CGI DNA methylation. Therefore, we propose KDM5C plays a fundamental role in the development of tissue identity during early embryogenesis, including the establishment of the soma-germline boundary. By systematically characterizing KDM5C’s role in germline gene repression, we unveiled divergent mechanisms governing the misexpression of distinct germline gene classes in somatic lineages.

By comparing *Kdm5c* mutant males and females, we revealed germline gene supression is sexually dimorphic. Female EpiLCs are more severely impacted by loss of KDM5C-mediated germline gene sup-pression, yet this difference is not due to the large number of germline genes on the X chromosome^54,55^. Heightened germline gene misexpression in females may be related to females having a higher dose of KDM5C than males, due to its escape from XCI^50–53^. Intriguingly, heterozygous knockout females (*Kdm5c^-/+^*) also had over double the number of germline DEGs than hemizygous knockout males (*Kdm5c^-/y^*), even though their expression of KDM5C should be roughly equivalent to that of wild-type males (*Kdm5c^+/y^*) . Males could partially compensate for KDM5C’s loss via the Y-chromosome homolog, KDM5D^14^. However, KDM5D has not been reported to regulate germline gene expression. Nevertheless, these results demonstrate germline gene silencing mechanims differ between males and females, which warrants further study to elucidate the biological ramifications and underlying mechanisms.

We found KDM5C is largely dispensable for promoting normal gene expression during development, yet is critical for suppressing ectopic developmental programs. While some germline genes, such as *Dazl*, are also expressed in the 2-cell stage, the inner cell mass, and naïve ESCs, they are silenced in epiblast stem cells/EpiLCs^6,42,47,72,73^. Our data suggest the 2-cell-like state reported in *Kdm5c*-KO ESCs^47^ likely reflects KDM5C’s primary role in germline gene repression (Figure 3). Germline gene misexpression in *Kdm5c*-KO EpiLCs may indicate they are differentiating into primordial germ cell-like cells (PGCLCs)^35,36,38^. Yet, *Kdm5c*-KO EpiLCs had normal cellular morphology and properly expressed markers for primed pluripotency, including *Otx2* which blocks EpiLC differentiation into PGCs/PGCLCs^74^. In addition to unimpaired EpiLC differentiation, *Kdm5c*-KO gross brain morphology is overall normal^21^ and hardly any brain-specific genes were significantly dysregulated in the amygdala and hippocampus (Figure 1). Thus, ectopic germline gene expression occurs in conjunction with overall proper somatic differentiation in *Kdm5c*-KO animals.

Our work provides novel insight into the cross-talk between H3K4me2/3 and CpGme, which are gen-erally mutually exclusive^75^. In EpiLCs, loss of KDM5C binding at a subset of germline gene promoters, e.g. *D1Pas1*, strongly impaired promoter CGI methylation and resulted in their long-lasting de-repression into adulthood. Removal of H3K4me2/3 at CGIs is a plausible mechanism for KDM5C-mediated germline gene suppression^22,56^, given H3K4me2/3 repell DNMT3 activity^67,68^. However, emerging work indicates many histone-modifying enzymes have non-cataltyic functions that influnce gene expression, sometimes even more potently than their catalytic roles^76,77^. Indeed, KDM5C’s catalytic activity was recently found to be dispensible for repressing *Dazl* in ESCs^47^. In our study, *Dazl*’s promoter still gained CpGme in *Kdm5c*-KO exEpiLCs, even with elevated H3K4me2. *Dazl* and a few other germline genes employ multiple repressive mechanisms to facilitate CpGme, such as DNMT3A/B recruitment via E2F6 and MGA^5,6,48,49^. Thus, while some germline CGIs require KDM5C-mediated H3K4me removal to overcome promoter CGI escape from CpGme^75,78^, others do not. These results also suggest the requirement for KDM5C’s catalytic activity can change depending upon the locus and developmental stage. Further experiments are required to determine if catalytically inactive KDM5C can suppress germline genes at later developmental stages.

By generating a comprehensive list of mouse germline-enriched genes, we revealed distinct derepressive mechanisms governing early versus late-stage germline programs. Previous work on germline gene silencing has focused on genes with promoter CGIs^7,75^, and indeed the majority of KDM5C targets in EpiLCs were germ cell identity genes harboring CGIs. However, over 70% of germline-enriched gene promoters lacked CGIs, including the many KDM5C-unbound germline genes that are de-repressed in *Kdm5c*-KO cells. CGI-free, KDM5C-unbound germline genes were primarily late-stage spermatogenesis genes and significantly enriched for RFX2 binding sites, a central regulator of spermiogenesis^65,66^. These data suggest that once activated during early embryogenesis, drivers of germline gene expression like *Rfx2*, *Stra8*, and *Dazl* turn on downstream germline programs, ultimately culminating in the expression of spermiogenesis genes in the adult *Kdm5c*-KO brain. Therefore, we propose KDM5C is recruited via promoter CGIs to act as a brake against runaway activation of germline-specific programs. Future studies should adress how KDM5C is targeted to CGIs.

The above work provides the mechanistic foundation for KDM5C-mediated repression of tissue and germline-specific genes. However, the contribution of these ectopic, tissue-specific genes towards neurolog-ical impairments is still unknown. In addition to germline genes, we also identified significant enrichment of muscle and liver-enriched transcripts within the *Kdm5c*-KO brain. Intriguingly, select liver and muscle-enriched DEGs do have known roles within the brain, such as the liver-enriched lipid metabolism gene *Apolipoprotein C-I (Apoc1)*^30^. *APOC1* dysregulation is implicated in Alzheimer’s disease in humans^79^ and overexpression of *Apoc1* in the mouse brain can impair learning and memory^80^. KDM5C may therefore be crucial for neurodevelopment by fine-tuning the expression of tissue-erniched, dosage-sensitive genes like *Apoc1*.

Given that germline genes have no known functions within the brain, their impact upon neruodevelopment is currently unknown. In *C. elegans*, somatic misexpression of germline genes via loss of *Retinoblastoma (Rb)* homologs results in enhanced piRNA signaling and ectopic P granule formation in neurons^81,82^. Ectopic testicular germline transcripts have also been observed in a variety of cancers, including brain turmors in *Drosophila* and mammals and shown to promote cancer progression^10,11,83–85^. Intriguingly, mouse models and human cells for other chromatin-linked NDDs also display impaired soma-germline demarcation^13,86,87^, such as mutations in DNA methyltransferase 3b (DNMT3B), H3K9me1/2 methyltransferases G9A/GLP, and methyl-CpG -binding protein 2 (MECP2). Recently, the transcription factor ZMYM2 (ZNF198), whose mutation causes a NDD (OMIM #619522), was also shown to repress germline genes by promoting H3K4me removal and CpGme^88^. Thus, KDM5C is among a growing cohort of neurodevelopmental disorders with erosion of the germline-soma boundary. Further research is required to determine the impact of these germline genes upon neuronal functions and the extent to which this phenomenon occurs in humans.

## Materials and Methods

### Classifying tissue-enriched and germline-enriched genes

Tissue-enriched differentially expressd genes (DEGs) were determined by their classification in a previ-ously published dataset from 17 male and female mouse tissues^25^. This study defined tissue expression as greather than 1 Fragments Per Kilobase of transcript per Million mapped read (FPKM) and tissue enrichment as at least 4-fold higher expression than any other tissue.

We curated a list of germline-enriched genes using an RNA-seq dataset from wild-type and germline-depleted (Kit^W/Wv^) male and female mouse embryos from embryonic day 12, 14, and 16^34^, as well as adult male testes^31^. Germline-enriched genes met the following criteria: 1) their expression is greater than 1 FPKM in wild-type germline 2) their expression in any wild-type somatic tissues^25^ does not exceed 20% of maximum expression in wild-type germline, and 3) their expression in the germ cell-depleted (Kit^W/Wv^) germline, for any sex or time point, does not exceed 20% of maximum expression in wild-type germline. We defined sperm and egg-biased genes as those whose expression in the opposite sex, at any time point, is no greater than 20% of the gene’s maximum expression in a given sex. Genes that did not meet this threshold for either sex were classified as ‘unbiased’.

### Cell culture

We utilized our previously established cultures of male wild-type and *Kdm5c* knockout (-KO) embryonic stem cells^43^. Sex was confirmed by genotyping *Uba1*/*Uba1y* on the X and Y chromo-somes with the following primers: 5’-TGGATGGTGTGGCCAATG-3’, 5’-CACCTGCACGTTGCCCTT-3’. Deletion of *Kdm5c* exons 11 and 12, which destabilize KDM5C protein^21^, was confirmed through the primers 5’-ATGCCCATATTAAGAGTCCCTG-3’, 5’-TCTGCCTTGATGGGACTGTT-3’, and 5’-GGTTCTCAACACTCACATAGTG-3’.

Emrbyonic stem cells (ESCs) and epiblast-like cells were cultured using previously established methods^39^. Briefly, ESCs were initially cultured in primed ESC (pESC) media consisting of KnockOut DMEM (Gibco#10829–018), fetal bovine serum (Gibco#A5209501), KnockOut serum replacement (Invitrogen#10828–028), Glutamax (Gibco#35050-061), Anti-Anti (Gibco#15240-062), MEM Non-essential amino acids (Gibco#11140-050), and beta-mercaptoethanol (Sigma#M7522). They were then transitioned into ground-state, “naïve” ESCs (nESCs) by culturing for four passages in N2B27 media containing DMEM/F12 (Gibco#11330–032), Neurobasal media (Gibco#21103–049), Gluamax (Gibco#35050-061), Anti-Anti (Gibco#15240-062), N2 supplement (Invitrogen#17502048), and B27 supplement without vitamin A (Invitrogen#12587-010), and beta-mercaptoethanol (Sigma#M7522). Both pESC and nESC media were supplemented with 3 *µ*M GSK3 inhibitor CHIR99021 (Sigma #SML1046-5MG), 1 *µ*M MEK inhibitor PD0325901 (Sigma #PZ0162-5MG), and 1,000 units/mL leukemia inhibitory factor (LIF, Millipore#ESG1107).

nESCs were differentiated into epiblast-like cells (EpiLCs, 48 hours) and extendend EpiLCs (exEpiLCs, 96 hours) by culturing in N2B27 media containing DMEM/F12, Neurobasal media, Gluamax, Anti-Anti, N2 supplement, B27 supplement (Invitrogen#17504044), and beta-mercaptoethanol supplemented with 10 ng/mL fibroblast growth factor 2 (FGF2, R&D Biotechne 233-FB), and 20 ng/mL activin A (R&D Biotechne 338AC050CF), as previously described^39^.

### Real time quantitative PCR (RT-qPCR)

nESCs were differentiated into EpiLCs as described above. Cells were lysed with Tri-reagent BD (Sigma #T3809) at 0, 24, and 48 hours of differentiation. RNA was phase separated with 0.1 *µ*L/*µ*L 1-bromo-3-chloropropane (Sigma #B9673) and then precipitated with with isopropanol (Sigma #I9516) and ethanol purified. For each sample, 2 *µ*g of RNA was reverse transcribed using the ProtoScript II Reverse transcriptase kit from New England Biolabs (NEB #M0368S) and primed with oligo dT. Expression of *Kdm5c* was detected using the primers 5’-CCCATGGAGGCCAGAGAATAAG-3’ 5’-CTCAGCGGATAAGAGAATTTGCTAC-3’ and normalized to TBP using the primers 5’-TTCAGAGGATGCTCTAGGGAAGA-3’ 5’-CTGTGGAGTAAGTCCTGTGCC-3’ with the Power SYBR™ Green PCR Master Mix (ThermoFisher #4367659).

### Western Blot

Total protein was extracted during nESC to EpiLC differentiation at 0, 24, and 48 hours by sonicating cells at 20% amplitude for 15 seconds in 2X SDS sample buffer, then boiling at 100°C for 10 minutes. Proteins were separated on a 7.5% SDS page gel, transferred overnight onto a fluorescent membrane, blotted for rabbit anti-KDM5C (in house, 1:500) and mouse anti-DAXX (Santa Cruz #(H-7): sc-8043, 1:500), and then imaged using the LiCor Odyssey CLx system. Band intensitity was quantified using ImageJ.

### RNA sequencing (RNA-seq) data analysis

After ensuring read quality via FastQC (v0.11.8), reads were then mapped to the mm10 *Mus musculus* genome (Gencode) using STAR (v2.5.3a), during which we removed duplicates and kept only uniquely mapped reads. Count files were generated by FeatureCounts (Subread v1.5.0), and BAM files were converted to bigwigs using deeptools (v3.1.3) and visualized by the UCSC genome browser^70^. RStudio (v3.6.0) was then used to analyze counts files by DESeq2 (v1.26.0)^26^ to identify differentially expressed genes (DEGs) with a q-value (p-adjusted via FDR/Benjamini–Hochberg correction) less than 0.1 and a log2 fold change greater than 0.5. For all DESeq2 analyses, log2 fold changes were calculated with lfcShrink using the ashr package^89^. MA-plots were generated by ggpubr (v0.6.0), and Eulerr diagrams were generated by eulerr (v6.1.1). Boxplots and scatterplots were generated by ggpubr (v0.6.0) and ggplot2 (v3.3.2). The Upset plot was generated via the package UpSetR (v1.4.0)^90^. Gene ontology (GO) analyses were performed by the R package enrichPlot (v1.16.2) using the biological processes setting and compareCluster.

### Chromatin immunoprecipitation followed by DNA sequecing (ChIP-seq) data analysis

ChIP-seq reads were aligned to mm10 using Bowtie1 (v1.1.2) allowing up to two mismatches. Only uniquely mapped reads were used for analysis. Peaks were called using MACS2 software (v2.2.9.1) using input BAM files for normalization, with filters for a q-value < 0.1 and a fold enrichment > 1. We removed “black-listed” genomic regions that often give aberrant signals. Common peak sets were obtained in R via DiffBind^91^ (v3.6.5). In the case of KDM5C ChIP-seq, *Kdm5c*-KO false-positive peaks were then removed from wild-type samples using bedtools (v2.25.0). Peak proximity to genomic loci was determined by ChIPSeeker^92^ (v1.32.1). Gene ontology (GO) analyses were performed by the R package enrichPlot (v1.16.2) using the biological processes setting and compareCluster. Enriched motifs were identified using HOMER^61^ to search for known motifs within 500 base pairs up and downstream of the transcription start site. Average binding across genes was visualized using deeptools (v3.1.3). Bigwigs were visualized using the UCSC genome browser^70^.

### CpG island (CGI) analysis

Locations of CpG islands were determined through the mm10 UCSC genome browser CpG island track^70^, which classified CGIs as regions that have greater than 50% GC content, are larger than 200 base pairs, and have a ratio of CG dinucleotides observed over the expected amount greater than 0.6. CGI genomic coordinates were then annotated using ChIPseeker^92^ (v1.32.1) and filtered for ones that lie within promoters of germline-enriched genes (TSS ± 500).

### Whole genome bisulfite sequencing (WGBS)

Genomic DNA (gDNA) from male naïve ESCs and extended EpiLCs was extracted using the Wizard Genomic DNA Purification Kit (Promega A1120), following the instructions for Tissue Culture Cells. gDNA from two wild-types and two *Kdm5c*-KOs of each cell type was sent to Novogene for WGBS using the Illumina NovaSeq X Plus platform and sequenced for 150 bp paired-end reads (PE150). All samples had greater than 99% bisulfite conversion rates. Reads were adapter and quality trimmed with Trim Galore (v0.6.10) and aligned to the mm10 genome using Bismark^93^ (v0.22.1). Analysis of differential methylation at gene promoters was perfomed using methylKit^71^ (v1.28.0) with a minimum coverage of 3 paired reads, a percentage greater than 25% or less than -25%, and q-value less than 0.01. methylKit was also used to calculate average percentage methylation at germline gene promoters. Methylation bedgraph tracks were generated via Bismark and visualized using the UCSC genome browser^70^.

### Data access

#### WGBS in wild-type and *Kdm5c*-KO ESCs and exEpiLCs

Raw fastq files are deposited in the Sequence Read Archive (SRA) https://www.ncbi.nlm.nih.gov/sra under the bioProject PRJNA1165148. https://www.ncbi.nlm.nih.gov/bioproject/PRJNA1165148

#### Published datasets

All published datasets are available at the Gene Expression Omnibus (GEO) https://www.ncbi.nlm.nih.gov/geo. Published RNA-seq datasets analyzed in this study included the male wild-type and *Kdm5c*-KO adult amygdala and hippocampus^24^, available at GEO: GSE127722. Male and female wild-type, *Kdm5c*-KO, and *Kdm5c*-HET EpiLCs^43^ are available at GEO: GSE96797.

Previously published ChIP-seq experiments included KDM5C binding in wild-type and *Kdm5c*-KO EpiLCs^43^ (available at GEO: GSE96797) and mouse primary neuron cultures (PNCs) from the cortex and hippocampus^21^ (available at GEO: GSE61036). ChIP-seq of histone 3 lysine 4 dimethylation (H3K4me2) in male wild-type and *Kdm5c*-KO EpiLCs^43^ is also available at GEO: GSE96797. ChIP-seq of histone 3 lysine 4 trimethylation (H3K4me3) in wild-type and *Kdm5c*-KO male amygdala^24^ are available at GEO: GSE127817.

#### Data analysis

Scripts used to generate the results, tables, and figures of this study are available via the GitHub repository: https://github.com/kbonefas/KDM5C_Germ_Mechanism

## Acknowledgements

We thank Drs. Sundeep Kalantry, Milan Samanta, and Rebecca Malcore for providing protocols and expertise in culturing mouse ESCs and EpiLCs, as well as providing the wild-type and *Kdm5c*-KO ESCs used in this study. We thank Dr. Jacob Mueller for his insight in germline gene regulation and directing us to the germline-depleted mouse models. We also thank Drs. Gabriel Corfas, Kenneth Kwan, Natalie Tronson, Michael Sutton, Stephanie Bielas, Donna Martin, and the members of the Iwase, Sutton, Bielas, and Martin labs for helpful discussions and critiques of the data. We thank members of the University of Michigan Reproductive Sciences Program for providing feedback throughout the development of this work. This work was supported by grants from the National Institutes of Health (NIH) National Institute of Neurological Disorders and Stroke (NS089896, 5R21NS104774, and NS116008 to S.I.), National institute of Mental Health (1R21MH135290 to S.I.), the Simons Foundation Autism Research Initiative (SFARI, SFI-AN-AR-Pilot-00005721 to S.I.), the Farrehi Family Foundation Grant (to S.I.), the University of Michigan Career Training in Reproductive Biology (NIH T32HD079342, to K.M.B.), the NIH Early Stage Training in the Neurosciences Training Grant (NIH T32NS076401 to K.M.B.), and the Michigan Predoctoral Training in Genetics Grant (NIH T32GM007544, to I.V.)

## Author Contributions

K.M.B. and S.I. conceived the study and designed the experiments. I.V. generated the ESC and exEpiLC WGBS data. K.M.B performed all data analysis and all other experiments. The manuscript was written by K.M.B and S.I. and edited by K.M.B, S.I., and I.V.

