## Supplementary Figures for "KDM5C is a sex-biased brake against germline gene expression programs in somatic lineages"

### Supplementary Tables

**Supplementary table 1: Misexpression of tissue-enriched genes within the *Kdm5c*-KO brain.** 1) DESeq2 results table of the *Kdm5c*-KO hippocampus, with tissue-enriched genes annotated 2) Same as 1 but for the *Kdm5c*-KO amygdala 3) Results of the Fisher exact test for enrichment tissue-specific genes. 4) enrichPlot gene ontology results of testis-enriched genes misexpressed in the amygdala or hippocampus.

**Supplementary table 2: Mouse germline-enriched genes.** List of mouse germline-enriched genes identified in this study (see Methods). Includes whether germline gene promoters have KDM5C ChIP-seq peaks and CpG islands.

**Supplementary table 3: Germline gene misexpression in *Kdm5c* mutant EpiLCs.** 1) DESeq2 results table of male *Kdm5c*-KO (*Kdm5c*$^{-/y}$) epiblast-like cells (EpiLCs), with annotations for tissue-enriched genes. 2) Results of Fisher exact test on XY EpiLC tissue-enriched genes. 3) enrichPlot gene ontology results of germline-enriched genes misexpressed in male *Kdm5c*-KO EpiLCs, amygdala, and hippocampus. 4) DESeq2 results table of XX *Kdm5c*-HET (*Kdm5c*$^{-/+}$) EpiLCs 5) DESeq2 results table of XX *Kdm5c*-KO (*Kdm5c*$^{-/-}$) EpiLCs 6) Germline genes misexpressed in XY *Kdm5c*-KO, XX *Kdm5c*-KO, and XX *Kdm5c*-HET EpiLCs. 7) enrichPlot gene ontology results of germline genes misexpressed in male versus female *Kdm5c* mutant EpiLCs

**Supplementary table 4: KDM5C ChIP-seq in EpiLCs and PNCs.** 1) enrichPlot gene ontology results of KDM5C-bound promoters (see Methods) in epiblast-like cells (EpiLCs) and forebrain primary neuron cultures (PNCs). 2) HOMER known motif analysis of germline gene promoters (TSS ± 500 bp) bound by KDM5C . 3) HOMER known motif analysis of germline genes not bound by KDM5C.

**Supplementary table 5: Germline gene CpG islands and promoter CpG methylation.** 1) enrichPlot gene ontology results of germline genes with and without CpG islands (CGIs) within their promoter (TSS ± 500 bp). 2) methylKit whole genome bisulfite sequencing (WGBS) results table comparing germline gene promoter CpG methylation (TSS ± 500 bp) in male wild-type ESCs versus wild-type extended EpiLCs (exEpiLCs). 3) methylKit WGBS results table comparing germline gene promoter CpG methylation in male *Kdm5c*-KO versus wild-type exEpiLCs. 4) enrichPlot gene ontology results of promoters hypomethylated in *Kdm5c*-KO exEpiLCs

### Supplementary Figures

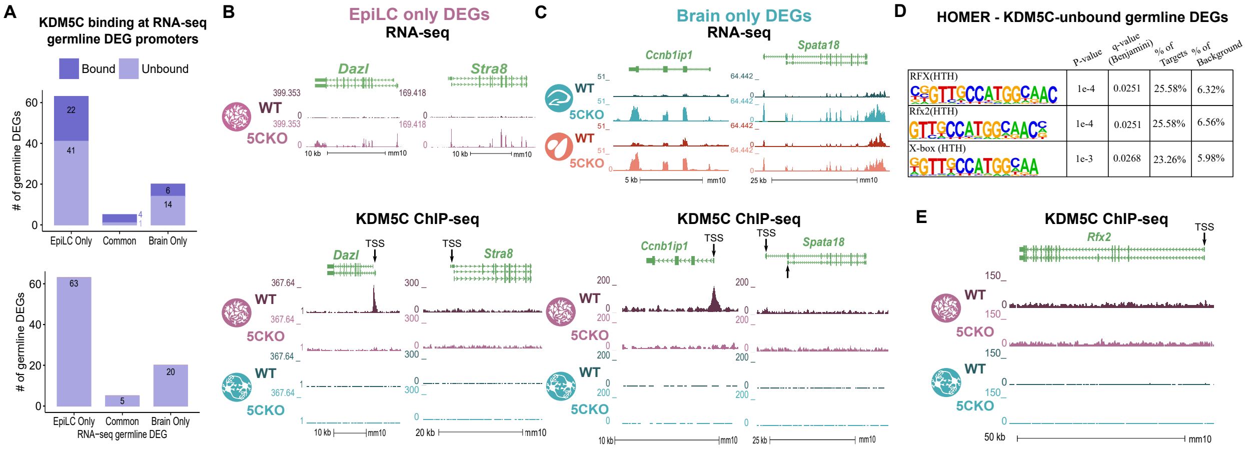


**Supplementary Figure 1: KDM5C binds to a subset of germline RNA-seq differentially expressed genes. A.** Bar graph of the number of germline-enriched DEGs with promoter KDM5C ChIP-seq peaks in wild-type EpiLCs (Top) and PNCs (Bottom). RNA-seq DEGs were classified as shared betwen EpiLCs and the brain (Common), unique to EpiLCs (EpiLC Only), or unique to one or multiple brain regions (Brain Only). **B.** Average bigwigs of two example RNA-seq DEGs dysregulated in EpiLCs but not the brain, *Dazl* and *Stra8*. Top is the RNA-seq tracks for wild-type (WT) and *Kdm5c*-KO (5CKO) EpiLCs, bottom is the KDM5C ChIP-seq tracks, with the annotated transcription start site (TSS) for each gene. **C.** Same as B but for two example DEGs only dysregualted in the brain and not expressed in EpiLCs, *Ccnb1ip1* and *Spata18*. **D.** HOMER motif analysis of all KDM5C-unbound germline DEGs shows signficant enrichment of multiple RFX members and their X-box motif. **E.** KDM5C ChIP-seq shows no KDM5C accumulation at the *Rfx2* promoter in EpiLCs or PNCs.

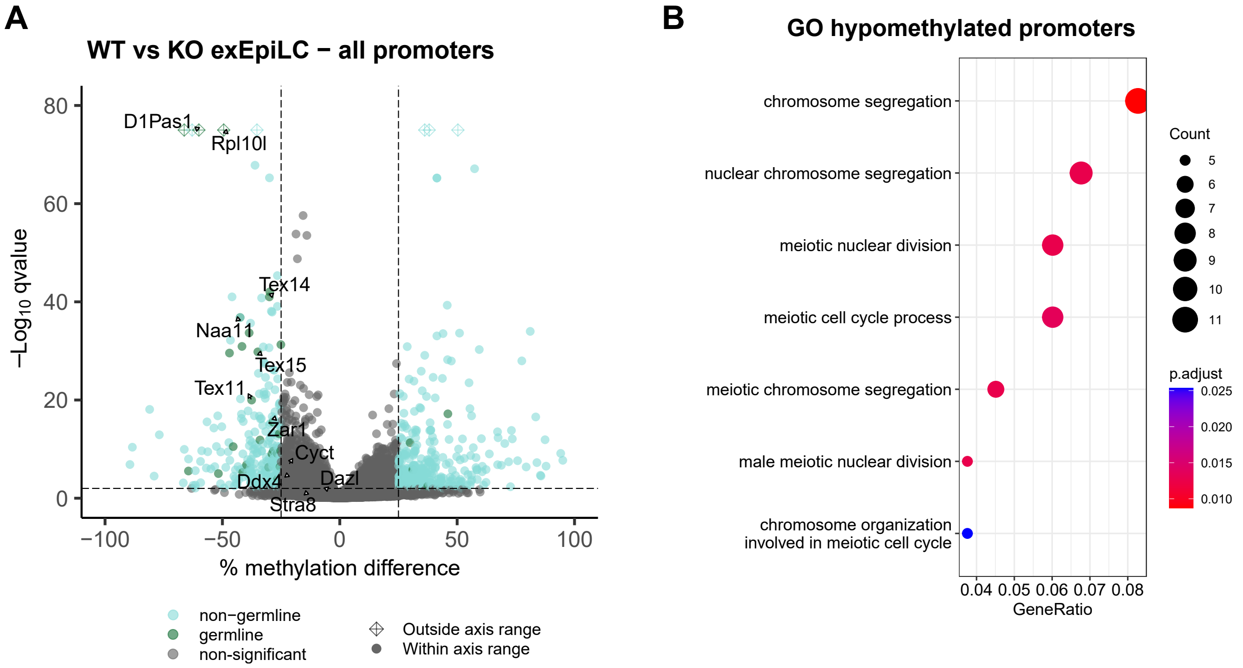


**Supplementary Figure 2: Loss of KDM5C impairs CpG methylation of germline gene promoters. A.** Volcano plot of whole genome bisulfite sequencing (WGBS) for all gene promoters in wild-type (WT) versus *Kdm5c*-KO extended EpiLCs (exEpiLCs). Significantly differentially methylated promoters ($q < 0.01$, |methylation difference| > 25%). Germline promoters highlighted in green, non-germline promoters in light blue, non-significant promoters in gray. **B.** enrichPlot gene ontology of all promoters significantly hypomethylated in *Kdm5c*-KO exEpiLCs.
